# Functional characterization of hypertriglyceridemia-associated mutations in Lipase Maturation Factor 1 by a second-generation activity assay

**DOI:** 10.64898/2026.09.03.749245

**Authors:** Candy Bedoya, Parmis Rejali, Xanden McCleary, Zachary Hall, Michael M. Hoffmann, Laurie Green, Philip H. Frost, Mary J. Malloy, Clive R. Pullinger, John P. Kane, Miklós Péterfy

## Abstract

Lipase maturation factor 1 (LMF1) is a chaperone for lipoprotein lipase (LPL) and critically required for the enzyme to attain lipase activity. LMF1 has been identified as a canonical gene affected in Familial Chylomicronemia Syndrome (FCS) and it has also been extensively analyzed in Multifactorial Chylomicronemia Syndrome (MCS), the polygenic form of severe hypertriglyceridemia (hTG). While recent genetic studies resulted in an explosion in the number of hTG-associated LMF1 variants in different populations, their functional significance remains largely unknown. Here, we present a second-generation LMF1 activity assay allowing the streamlined quantitative functional analysis of LMF1 variants. The assay is based on the reconstitution of lipase maturation in transfected LMF1-deficient cells and fluorescence-based measurement of LPL activity in the culture medium. The use of Gaussia luciferase (GLuc)-LMF1 fusion constructs allows the assessment of LMF1 protein expression and the calculation of LMF1 specific activity. Several variants previously reported as neutral resulted in loss of LMF1 function in the new assay demonstrating increased sensitivity to detect variants of modest effect sizes. Furthermore, the functional analysis of 14 hTG-associated missense mutations identified 12 loss-of-function (LOF) variants. To gain initial insights into the structural determinants of LMF1 function, we analyzed the domain distribution of 73 hTG-associated missense variants and identified Loop C as a critical region in lipase maturation. In conclusion, we developed a simple, non-radioactive and sensitive quantitative assay of LMF1 activity to facilitate the functional analysis of hyperlipidemia-associated genetic variants and structure-function studies.

## INTRODUCTION

Lipase maturation factor 1 (LMF1) is an endoplasmic reticulum (ER) membrane-localized chaperone first identified as the protein affected by the *combined lipase deficiency* (*cld*) mutation in a naturally occurring mouse model of chylomicronemia and severe hypertriglyceridemia (1). The underlying pathophysiological defect in *cld* mice is impaired catabolism of TG-rich lipoproteins due to combined lipoprotein lipase (LPL) and hepatic lipase (HL) deficiency. Biochemical studies revealed that LMF1 is required for the post-translational maturation of nascent LPL, and to a lower extent HL, polypeptides into active enzymes within the lumen of the ER. Subsequently, LMF1 has been identified as a redox chaperone facilitating the formation of disulfide bonds, which are critical for LPL folding and enzymatic activity (2).

Following its identification in the mouse, LMF1 has become a candidate gene in human hypertriglyceridemia. Indeed, genetic analyses identified multiple homozygous or biallelic mutations in patients with Familial Chylomicronemia Syndrome (FCS), a rare monogenic form of severe hypertriglyceridemia, and LMF1 is now considered one of five canonical genes (the others being LPL, APOC2, APOA5 and GPIHBP1) in this disease (3). Furthermore, rare heterozygous loss-of-function variants in LMF1 have been identified in Multifactorial Chylomicronemia Syndrome (MCS), a more common polygenic manifestation of severe hypertriglyceridemia (4). Interestingly, genome-wide association studies also revealed common variants at the LMF1 locus to be associated with TG levels in the general population (5, 6). Collectively, these results indicate that LMF1 gene variants across a diverse allelic spectrum contribute to pathological and normal variation in TG levels in the population.

As a canonical gene in FCS, the LMF1 gene is now routinely sequenced in pursuit of a genetic diagnosis in severe hypertriglyceridemia and over 100 rare coding variants have been reported (4, 7–13). However, only a handful of these variants have been functionally evaluated and most remain classified as variants of uncertain significance. Consequently, clinical insights from genotyping results are limited and there is a need for the experimental evaluation of LMF1 activity to assess the pathogenicity of mutations associated with hypertriglyceridemia.

We have previously developed an *in vitro* LMF1 activity assay (from here on referred to as ‘first-generation assay’) based on the reconstitution of LPL-maturation activity in transfected LMF1-deficient cells (14). In this assay, *cld*-mutant cells are co-transfected with LMF1, LPL and secreted alkaline phosphatase (SEAP) expression vectors followed by the measurement of LPL and SEAP activities secreted into the culture media. LPL activity serves as a quantitative proxy for LMF1 function, whereas SEAP is used to control for variation in transfection efficiency. This assay has been used for the functional characterization of several hypertriglyceridemia-associated LMF1 variants in different populations (1, 11, 15, 16).

Although the first-generation assay has proven useful for the evaluation of LMF1 variants of large impact, as typically observed in FCS, it has limitations in the analysis of the broad spectrum of allelic effects associated with missense variants occurring in MCS. First, as medium LPL activity is a composite readout of LMF1 activity and expression level, changes in protein turnover confounds the interpretation of variants’ true effect on LMF1 activity. Indeed, several previously identified variants result in significantly impaired LPL maturation that is due in part, or even entirely, to reduced LMF1 expression (14, 16, 17). Thus, it is critical that the functional evaluation of LMF1 variants includes the assessment of protein expression and be based on the determination of LMF1 specific activity (i.e. LPL activity/LMF1 protein mass). While Western blotting has been used to evaluate the impact of LMF1 variants on protein expression, the semiquantitative and labor-intensive nature of this technique limits the precision and throughput of the assay.

Additional limitations of the first-generation assay include low throughput owing to a multi-step LPL activity assay based on radiolabeled triolein substrate and the reliance on mouse *cld* hepatocytes for complementation. The latter results in low assay sensitivity due to the relatively poor transfectability of these cells and a species mismatch when analyzing human LMF1 variants. Finally, while the secreted nature of SEAP makes it a convenient marker to assay, it is not well-suited for transfection normalization, because its activity is affected by ER stress (18), a process that may be influenced by LMF1 expression. Indeed, it has been proposed that LMF1 plays a general role in ER homeostasis beyond lipase maturation and affects the oxidative folding of many other proteins (2). The objective of the present work is to address these limitations and develop an improved second-generation assay for LMF1 lipase maturation activity.

## MATERIALS AND METHODS

### Reagents

Cell culture supplies and the EnzChek fluorescent lipase substrate (#E33955) was obtained from ThermoFisher, Inc. All other reagents, including the Zwitterionic detergent SB3-14 (Zwittergent 3-14, #693017) and branched polyethylenimine (PEI, #408727) were obtained from Sigma-Aldrich, Inc.

### Fluorescent LPL activity assay

*Overview*. LPL activity in cell culture media was assessed using a fluorescent lipase assay based on published protocols with modifications (19, 20). A 200 μM stock solution of EnzChek fluorescent substrate in DMSO was freshly diluted in 2x assay buffer to obtain the EnzChek Working Reagent. Fifty μl per well of purified LPL or LPL-containing media was added to a black 96-well plate and preincubated at 37 °C for 15 min. After the addition of 50 μl of prewarmed Working Reagent, the plate was immediately read in a SpectraMax M2e plate reader (Molecular Devices) for kinetic (1-minute intervals for 10-15 min) fluorescence (482 nm excitation, 515 nm emission, 495 nm filter cutoff) measurement. LPL activity was estimated from the slope of the linear portion of fluorescence curve (typically between 30-360 sec), as determined by curve fitting with the SoftMax Pro7 software.

*Assay optimization*. As assay buffers of different compositions have been reported in EnzChek assays, we determined optimal buffer conditions through a side-by-side comparison of several buffer formulations. Purified LPL (Part #261004-T, Cell Biolabs, Inc.) at 50 mU/ml diluted in cell culture media (DMEM, 10% FBS) was assayed in duplicates in the presence of 8 μM EnzChek in either Buffer A (Part #261002, Cell Biolabs, Inc.), Buffer B (19), Buffer C (Buffer B with 0.15 M NaCl), Buffer D (20) or Buffer E (Buffer D with 2 mM CaCl2). To assess the effect of substrate concentration on LPL activity, 25 mU/ml LPL was diluted in cell culture medium (DMEM, 10% FBS) and assayed in the presence of 0, 3, 5, 7 or 14 μM EnzChek in Buffer B. The linear range of the assay was evaluated by determining the initial rate of fluorescence increase in the presence of 0, 31, 62 and 125 mU/ml purified LPL and 7 μM EnzChek in Buffer B.

### Generation of LMF1-deficient cell lines

A double-stranded DNA fragment corresponding to the gRNA targeting sequence of ACATCCCGGATTGTCCTGTG within exon 4 of the LMF1 gene was cloned into the BbsI site of the PX495 vector (#62988, Addgene). Transfected HEK293FT cells (R70007, ThermoFisher) were selected with 2 μg/ml puromycin for 7 days followed by a 7-day expansion in the absence of selection. Cells were plated at limiting dilution and 20 colonies were grown to clonal cell lines. The success of CRISPR mutagenesis was assessed by the amplification of exon 4 using primers GACTAAGCTTGGATGGGAGTCCCAGCTTCTGGA and CACTGAGCTCCACGCAGAAGAGCTCCACT and separation of PCR products by 2.5% agarose gel electrophoresis. Nine clones lacking a wild-type-sized DNA fragment were selected for the functional analysis of LMF1 deficiency. Cells were co-transfected with or without pcDNA3.1-LMF1 and a mixture of the pcDNA6-LPL (14) and pCMV-GLuc (New England Biolabs) expression vectors followed by the analysis of LPL activity in culture media and Gaussia luciferase (GLuc) activity in cell lysates to confirm successful transfection. Mutations in functionally LMF1-deficient cell lines were identified by cloning and sequencing of PCR products containing exon 4. A HEK293FT clonal cell line (#13) with a homozygous 7-bp deletion and premature stop codon in exon 4 was used for LMF1 activity assays.

### Generation of expression constructs

#### GLuc-LMF1 plasmid

For initial experiments, we generated pcDNA3.1-GLuc-LMF1 expression vectors as follows. GLuc was PCR-amplified without its signal sequence and termination codon with primers GGCC<u>GGATCC</u>ACCATGAAGCCCACCGAGAACAACGA and GCTCGA <u>GCGGCCGC</u>GTCACCACCGGCCCCCTT using the pCMV-GLuc vector as template and cloned between the BamHI/NotI sites in pcDNA3.1. Mutant LMF1 cDNA sequences were generated by site-directed mutagenesis, as previously described (15), PCR-amplified with primers GACTCA<u>GCGGCCGC</u>TCGCCCTGACAGCCCAA and CGTCATC<u>GTTTAAAC</u>TAGAGGGGCCCGGGCAGAGGCCA and cloned between the NotI/PmeI sites in pcDNA3.1-GLuc.

#### GLuc-LMF1/FLuc plasmid

In initial experiments, the pcDNA3.1-GLuc-LMF1 vectors were co-transfected with a separate plasmid expressing firefly luciferase (FLuc) to control for variability in transfection efficiency. To improve assay normalization and streamline the experimental protocol, we incorporated the FLuc gene into the pcDNA3.1-GLuc-LMF1 plasmid as follows. A ∼2.2-kb restriction fragment containing FLuc/SV40-pA/SV40-enhancer was obtained by HindIII/BamHI digests of the pGL3-Control plasmid (Promega) and used to replace the ∼1.2-kb AvrII/BstZ17I fragment containing the Neomycin/SV40-pA cassette in pcDNA3.1-GLuc-LMF1 after blunting. The resulting pcDNA3.1-GLuc-LMF1/FLuc plasmid contains two separate transcriptional units driven by the CMV (GLuc-LMF1 gene) and SV40 (FLuc gene) promoters and was used in the LMF1 activity assays.

### LMF1 activity assay

#### Experimental procedure

HEK293FT-LMF1-KO (clone #13) cells were seeded in 48-well plates and co-transfected in quintuplicate wells with a mixture of 0.25 ng/well pcDNA3.1-GLuc-LMF1/FLuc (or pcDNA3.1-GLuc/FLuc as negative control), 500 ng/well pcDNA6-LPL and 1 μl/well PEI in Opti-MEM for 6 hours followed by replacement with complete media (DMEM/10% FBS). One day after transfection, cell-associated LPL was removed by a 30-min preincubation with 10 U/ml heparin in complete media followed by replacement with 125 μl/well fresh heparin-containing media. Six hours later, culture media were collected for LPL assay and cells were lysed in 80 μl/well Passive Lysis Buffer (#E1941, Promega) by gentle shaking for 15 min followed by 3 rounds of freeze/thaw cycles. The latter procedure resulted in ∼2-fold increased GLuc activity and reduced replicate variability, whereas FLuc activity remained unaffected (data not shown). FLuc and GLuc activities in cell lysates were assessed with the Dual-Luciferase Reporter Assay System (#E1980, Promega) in a GloMax 96 dual-injector luminometer (Promega).

#### Calculations

LPL activity in each well was corrected by subtracting the average activity measured in control wells transfected with a vector lacking LMF1 (pcDNA3.1-GLuc/FLuc). FLuc and GLuc activities in each well were corrected by subtracting the average activities observed in mock-transfected cell lysates. Total LMF1 activity was defined as LPL activity in medium after normalization by FLuc activity in cell lysates to control for variation in transfection efficiency. LMF1 specific activity was defined as the ratio of medium LPL activity and cellular LMF1 protein mass, as determined by GLuc activity assays (i.e. LPL activity/GLuc activity). In each experiment, quintuplicate replicates were averaged and the results are presented as the mean ± SEM of at least three independent experiments.

### Study subjects

Hypertriglyceridemic subjects with fasting triglyceride levels above 400 and 885 mg/dl were identified in the Genomic Resource in Arteriosclerosis and Metabolic Disease at the Cardiovascular Research Institute, University of California, San Francisco (UCSF). LMF1 coding regions and exon-intron boundaries were sequenced as described (17). All subjects gave informed consent and the study was approved by the respective institutional review boards.

### Variant-effect prediction

Scores assigned to LMF1 variants by variant-effect prediction algorithms were obtained from dbNSFv4 (21).

### Statistical analyses

Results are presented as means ± SEM and analyzed by two-tailed unpaired t-test followed by the calculation of Benjamini-Hochberg False Discovery Rate. P values were adjusted for multiple comparisons (Holm-Sidak method) with GraphPad Prism 11.0.2. Group comparisons with adjusted p values < 0.05 were considered statistically significant. Binary classification was evaluated using Receiver Operating Characteristic (ROC) and Matthews Correlation Coefficient (MCC) analyses after threshold optimization with ThresholdTuner (22).

## RESULTS

### Development of a second-generation LMF1 activity assay

#### Assay overview

As schematically shown in **Figure 1**, the basic principle and overall workflow of the second-generation assay are similar to the previously established assay (14). LMF1-deficient cells are co-transfected with LMF1 and LPL expression vectors followed by the determination of LPL activity in culture media, which serves as a proxy for cellular LMF1 activity. However, several improvements have been implemented in the new assay. First, the LMF1 expression vectors include Gaussia luciferase (GLuc)-LMF1 fusion constructs, which allows precise quantitation of LMF1 protein expression and enables the calculation of LMF1 specific activity (LPL activity in media/GLuc activity in cell lysate). For transfection normalization, we replaced secreted alkaline phosphatase (SEAP) with a cytoplasmic enzyme, Firefly luciferase (FLuc). This change mitigates the potential confounding effects of altered ER redox state on SEAP activity (18) due to variable expression of LMF1, a redox chaperone (2). Moreover, the FLuc gene has been incorporated into the GLuc-LMF1 expression vector, as opposed to being delivered on a separate plasmid, to ensure identical LMF1:FLuc DNA ratios in all LMF1 test constructs. Finally, mouse *cld*-mutant hepatocytes have been replaced by highly transfectable LMF1-deficient HEK293FT (LMF1-KO) cells to increase assay sensitivity and ensure compatibility with human LMF1 test constructs.

**Figure 1.**
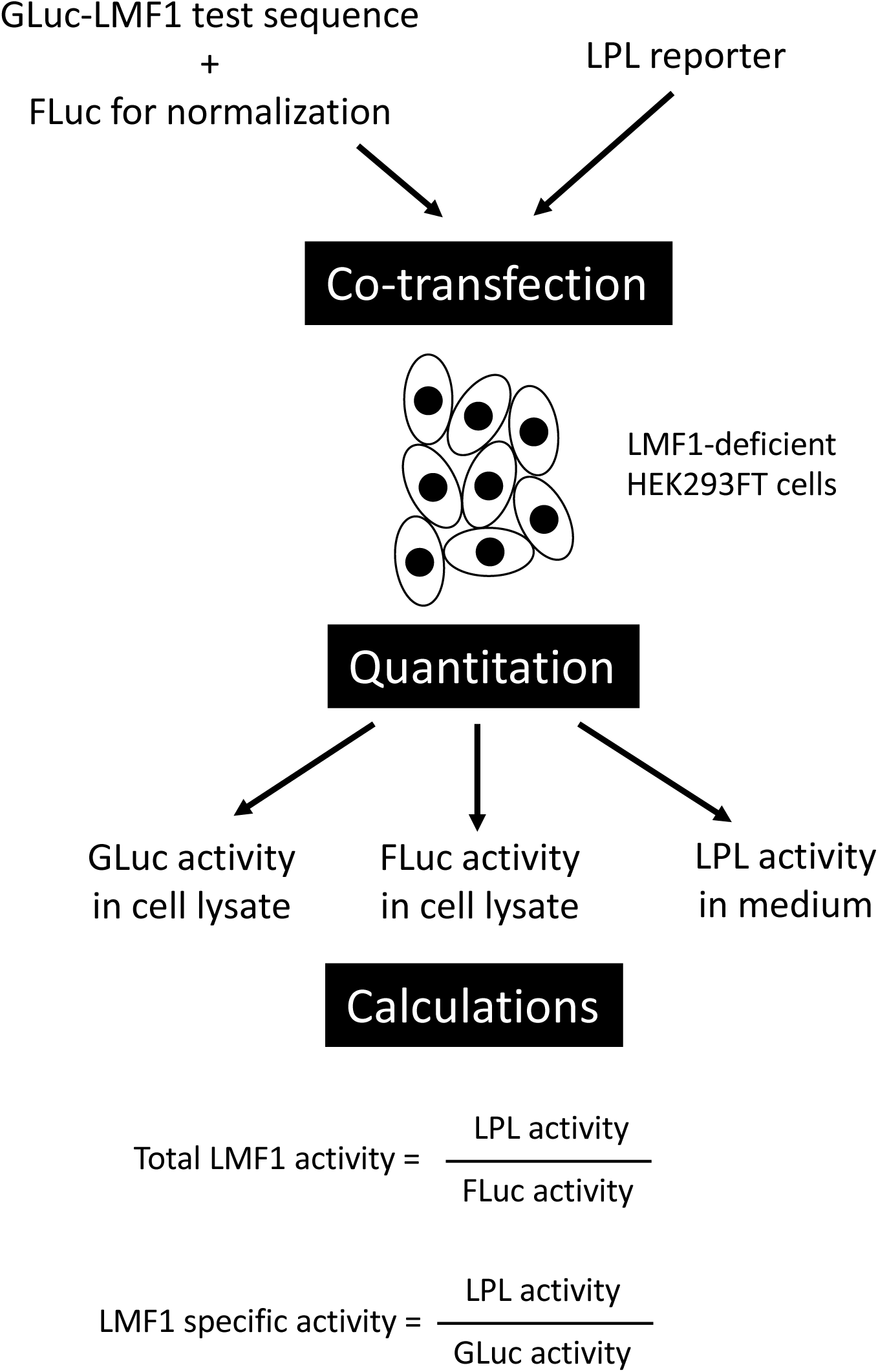
Overview of the second-generation LMF1 activity assay. LMF1-deficient HEK293FT cells are co-transfected by two vectors, one expressing the Gaussia luciferase (GLuc)-LMF1 test constructs and Firefly luciferase (FLuc), and the other LPL. GLuc serves as a proxy for LMF1 protein mass and FLuc is used for normalization. A day after transfection cell culture medium is harvested for LPL activity measurement and cells are lysed for dual luciferase (FLuc+GLuc) assay. FLuc-normalized LPL activity and LMF1 specific activity (LPL activity/GLuc activity) are reported.

#### Optimization of fluorescence-based LPL activity assay

In the first-generation assay, LPL activity was measured using radiometry requiring the preparation of lecithin-stabilized radiolabeled triolein substrate, fatty acid extraction and scintillation counting (23). To streamline this procedure and increase the throughput of the assay, we adapted a non-radioactive 96-well LPL activity assay based on the fluorescent EnzChek lipase-substrate and SB3-14, a zwitterionic detergent to solubilize the substrate (19, 20). Using purified LPL, we first compared commercial and published assay buffer compositions with different concentrations of NaCl, CaCl_2_ and BSA. Highest LPL activity was observed in Buffer B (**Fig. 2A**), which was used in subsequent assays.

**Figure 2.**
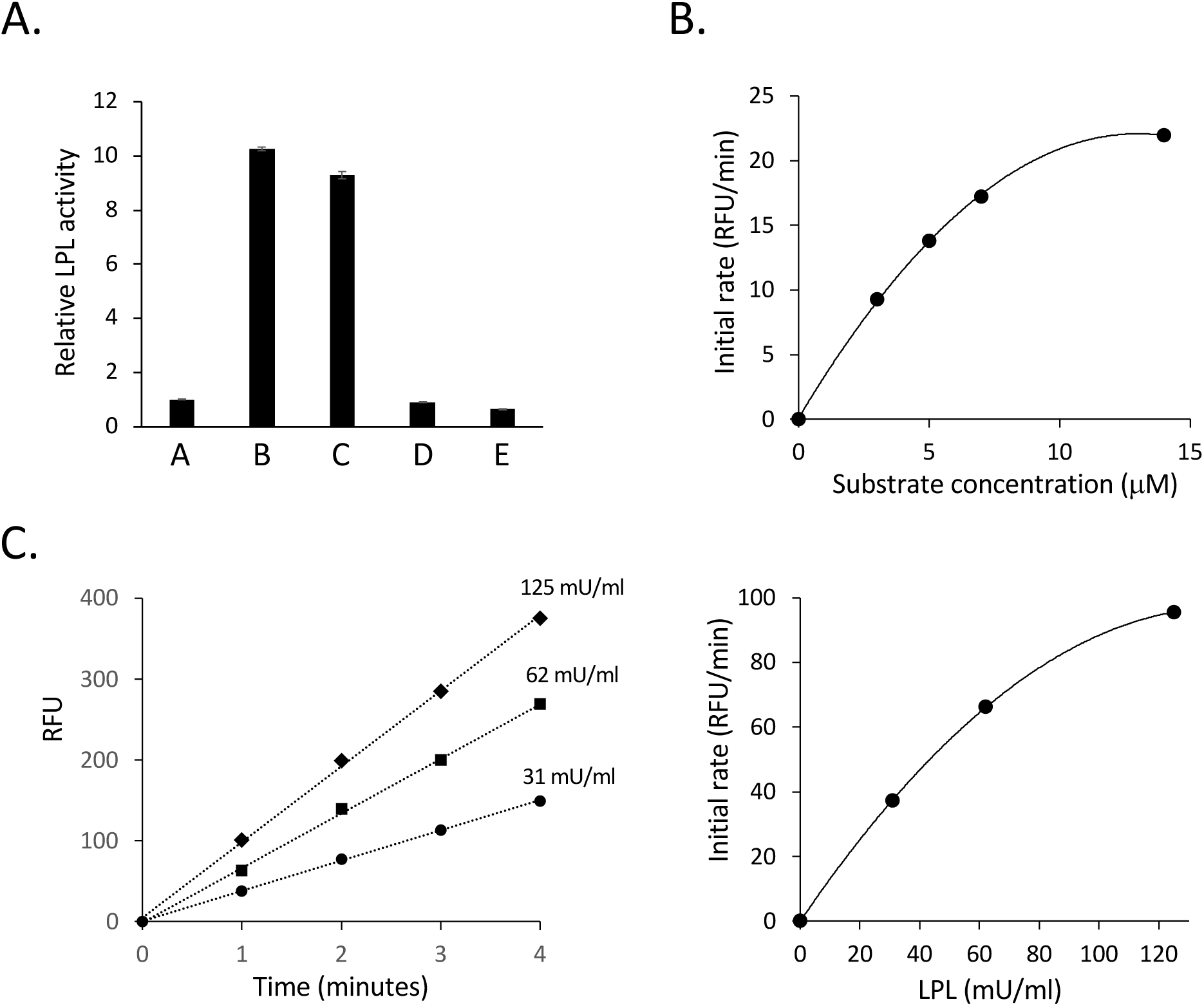
Optimization of fluorescent LPL assay. (**A**) Relative LPL activity (50 mU/ml) assayed in different buffer formulations A-E (see Methods for buffer compositions) in the presence of 8 μM EnzChek substrate. A commercially available LPL assay buffer (A) was used as reference. (**B**) The effect of EnzChek substrate concentration on the initial rate of fluorescence increase was determined with 25 mU/ml LPL in buffer B. (**C**) Testing the linear range of the LPL assay in buffer B at 7 μM EnzChek concentration. Panel on the left shows the initial increase of relative fluorescence (RFU) during the first 4 minutes of the assay at LPL concentrations of 31, 62 and 125 mU/ml. Rates of fluorescence increase are plotted as a function of LPL concentration on the right panel.

To find the optimal substrate concentration in the assay, we used a fixed amount of LPL and increasing amounts of EnzChek. LPL activity increased nearly linearly up to ∼7 μM substrate and plateaued at 14 μM (**Fig. 2B**). The concentration range in which the initial rate of fluorescence increase is proportional to LPL activity was determined by assaying increasing amounts of LPL at a fixed substrate concentration of 7 μM. The rate of fluorescence increase changed nearly linearly with LPL activity up to ∼60 mU/ml and plateaued at ∼120 mU/ml (**Fig. 2C**). Taken together, we established an optimized fluorescence-based LPL activity assay using 7 μM EnzChek substrate in a buffer containing 50 mM Tris-HCl, pH 8.0; 0.1 mg/ml SB3-14; 1 mg/ml fatty acid-free BSA; 2 mM CaCl_2_ and a linear range of up to ∼60 mU/ml LPL activity.

#### Generation of LMF1-deficient cell lines

The first-generation LMF1 activity assay is based on the reconstitution of LMF1 expression in a mouse hepatocyte cell line harboring the naturally occurring *combined lipase deficiency* (*cld*) mutation at homozygosity (14). The use of *cld* cell line in this assay has limitations including relatively poor transfectability and the expression of a truncated form of LMF1, which may retain some functionality (1). Thus, we used CRISPR-mediated gene targeting to generate LMF1-deficient HEK293FT cells. To assess functional LMF1 deficiency in clonal cell lines, we evaluated the effect of LMF1 reconstitution on LPL maturation in transient transfection assays. Overexpression of LMF1 in unmanipulated HEK293FT cells had no effect on LPL activity secreted into the culture media (WT in **Fig. 3A**). In contrast, transfection of LMF1 in several CRISPR-targeted cell lines resulted in 7-12-fold increases in LPL activity demonstrating functional LMF1 deficiency. Sequencing of the targeted LMF1 exon revealed a homozygous frame-shifting deletion in cell line #13, which was selected for the development of LMF1 activity assay.

**Figure 3.**
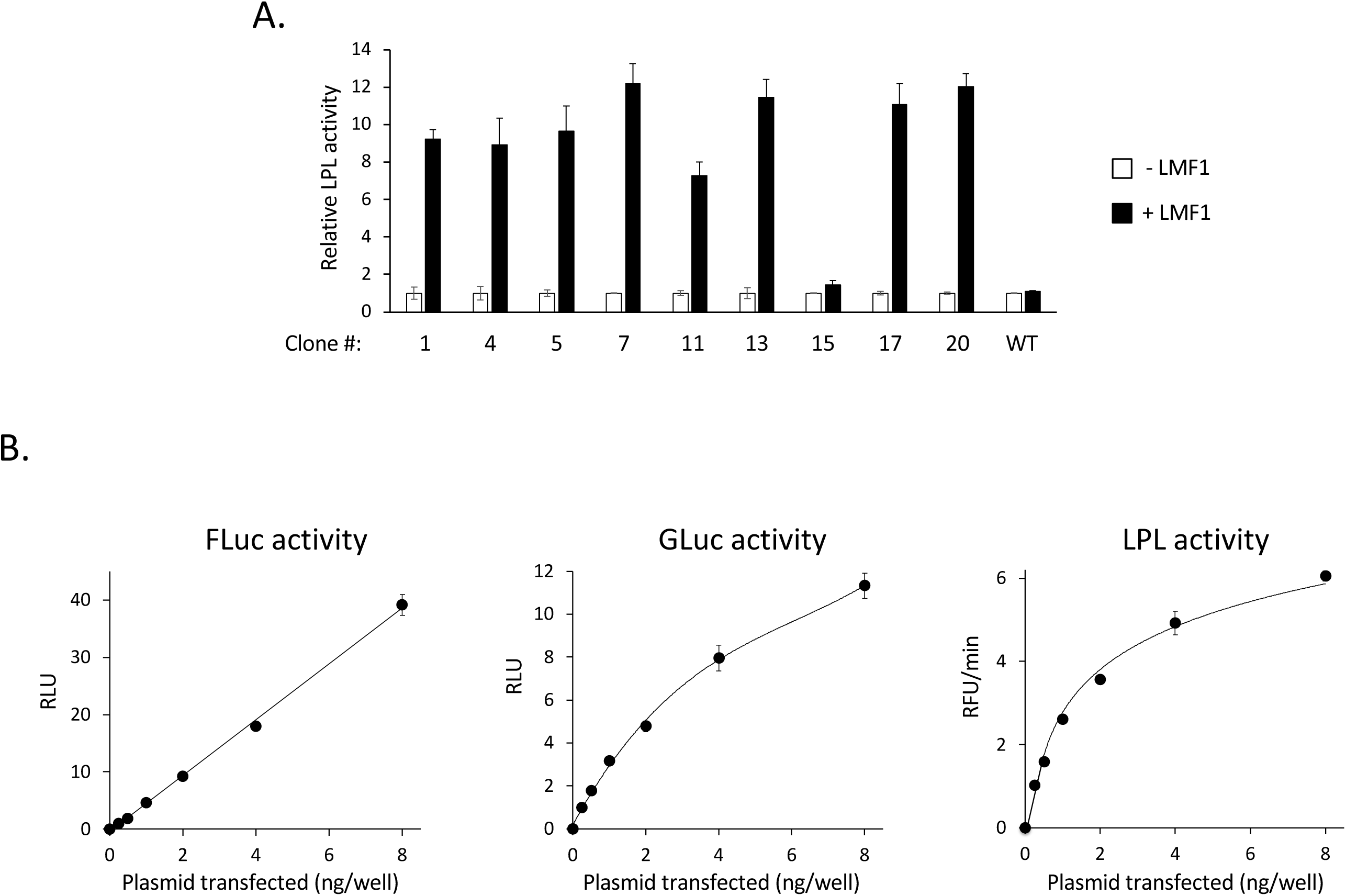
Development of the LMF1 activity assay. (**A**) Relative LPL activity in the cell culture media of LMF1 CRISPR-targeted HEK293FT clonal cell lines transiently transfected with or without wild-type LMF1. Untargeted (WT) HEK293FT cell line serves as control. (**B**) Cell-associated FLuc, GLuc and media LPL activity in HEK293FT clone #13 cells transfected with different amounts of LMF1 plasmid.

#### Assessment of the linear range of the assay

To establish the linear range of the LMF1 assay, we co-transfected variable amounts of GLuc-LMF1/FLuc test constructs (0.25-8 ng per well) together with a constant amount of LPL plasmid (500 ng per well) and determined the activities of all three reporters. While FLuc activity showed a linear response across the entire DNA concentration range, GLuc and LPL activities started to level off above ∼1 and ∼0.5 ng per well, respectively (**Fig. 3B**). We conclude that the assay is linear up to 0.5 ng LMF1 plasmid per 48-well and used 0.25 ng plasmid per well in subsequent assays.

### Functional assessment of LMF1 variants associated with hypertriglyceridemia

#### Reevaluation of previously characterized variants

To illustrate the utility and improved sensitivity of the second-generation LMF1 assay, we reanalyzed 8 hypertriglyceridemia-associated missense variants previously reported having no effect on LMF1 activity based on the first-generation assay (11, 15). While 3 variants were confirmed as neutral, 5 variants significantly reduced total LMF1 activity (i.e. normalized media LPL activity) in the new assay (**Table 1**). Furthermore, the analysis of LMF1 specific activity revealed that 7 of the 8 variants impaired the lipase-maturation function of the protein. We also assayed T143M, a variant that was previously reported as ‘gain-of-function’ (13). In the present assay, T143M significantly reduced total, but not specific, LMF1 activity indicating that this variant affects lipase maturation at least in part by decreasing LMF1 protein expression (**Table 1**). Importantly, all loss-of-function (LOF) variants identified in these analyses retained substantial (50-80%) LMF1 activity indicating the improved sensitivity of the second-generation assay to detect LOF variants with modest effect sizes.

**TABLE 1.** Functional analysis of HTG-associated LMF1 variants.

| LMF1 variant | Reference | Total LMF1 activity<sup>a</sup> | P value<sup>b</sup> | q value<sup>c</sup> | LMF1 sp. activity<sup>a</sup> | P value<sup>b</sup> | q value<sup>c</sup> |
| --- | --- | --- | --- | --- | --- | --- | --- |
| Previously reported as neutral: |  |  |  |  |  |  |  |
| R230Q | Surendran et al. (15) | 84 ± 7 | 0.094 | 0.103 | 72 ± 6 | 0.016 | **0.026** |
| R264C | Surendran et al. (15) | 88 ± 6 | 0.153 | 0.160 | 78 ± 5 | 0.031 | **0.045** |
| R351Q | Surendran et al. (15) | 77 ± 7 | 0.044 | 0.053 | 77 ± 7 | 0.049 | 0.056 |
| R354W | Surendran et al. (15) | 70 ± 7 | 0.029 | **0.039** | 72 ± 7 | 0.040 | **0.048** |
| R364Q | Surendran et al. (15) | 49 ± 3 | <0.001 | **<0.001** | 50 ± 3 | <0.001 | **<0.001** |
| R451W | Surendran et al. (15) | 73 ± 5 | 0.015 | **0.023** | 72 ± 7 | 0.027 | **0.041** |
| R523H | Surendran et al. (15) | 74 ± 5 | 0.011 | **0.021** | 76 ± 5 | 0.014 | **0.026** |
| P562R | Plengpanich et al. (11) | 80 ± 4 | 0.013 | **0.021** | 65 ± 2 | 0.003 | **0.008** |
| Previously reported as gain-of-function: |  |  |  |  |  |  |  |
| T143M | Dancer et al. | 43 ± 9 | 0.005 | **0.012** | 79 ± 18 | 0.319 | 0.319 |
| Previously uncharacterized HTG-associated variants: |  |  |  |  |  |  |  |
| R216T | present study | 26 ± 2 | <0.001 | **<0.001** | 23 ± 2 | <0.001 | **<0.001** |
| G228E | Dron et al. (4) | 62 ± 3 | <0.001 | **0.003** | 57 ± 9 | 0.015 | **0.026** |
| G228A | Dron et al. (4) | 58 ± 4 | 0.001 | **0.003** | 62 ± 6 | 0.006 | **0.014** |
| M238T | Dron et al. (4) | 77 ± 5 | 0.024 | **0.035** | 68 ± 8 | 0.038 | **0.048** |
| G284S | Dron et al. (4) | 37 ± 4 | <0.001 | **<0.001** | 28 ± 5 | <0.001 | **0.001** |
| G360S | Gill et al. (10) | 71 ± 3 | 0.001 | **0.003** | 60 ± 8 | 0.015 | **0.026** |
| V382M | Dron et al. (4) | 103 ± 5 | 0.351 | 0.351 | 88 ± 3 | 0.036 | **0.048** |
| A431D | Dancer et al. (13) | 72 ± 8 | 0.036 | **0.046** | 76 ± 9 | 0.059 | 0.065 |
| P457L | present study | 54 ± 8 | 0.010 | **0.021** | 60 ± 2 | <0.001 | **0.001** |
| R461C | Abedi et al. (7) | 4.7 ± 1 | <0.001 | **<0.001** | 3.6 ± 1 | <0.001 | **0.002** |
| R513Q | present study | 13 ± 3 | <0.001 | **<0.001** | 13 ± 5 | <0.001 | **0.001** |
| S522I | Johansen et al. (31) | 53 ± 5 | 0.002 | **0.005** | 48 ± 1 | <0.001 | **<0.001** |
| E531D | Gill et al. (10) | 84 ± 7 | 0.081 | 0.093 | 93 ± 10 | 0.313 | 0.319 |
| G532S | Gill et al. (10) | 80 ± 3 | 0.013 | **0.021** | 58 ± 2 | 0.001 | **0.003** |

<sup>a</sup> Values are expressed as % of wild-type LMF1 (Mean ± SEM).<sup>b</sup> Student's t-test P values (vs wild-type LMF1)<sup>c</sup> False Discovery Rate q values <0.05 appear in bold.

#### Functional analysis of previously untested variants

Next, we analyzed 14 LMF1 variants identified in patients with severe hypertriglyceridemia (**Table 1**). Twelve variants reduced total and/or specific LMF1 activity with a wide range of residual activities from 4-88% of wild-type protein. The R461C variant nearly completely (>95%) abolished LMF1 activity and identifies a residue critical for LMF1 function.

### Domain-distribution of hypertriglyceridemia-associated LMF1 variants

LMF1 is a multipass membrane-spanning protein consisting of 5 transmembrane (TM1-5) and 6 soluble (N-terminal, loops A-D, C-terminal) regions (**Figure 4A**). To gain further insights into the structure-function relationship in LMF1, we set out to determine the distribution of hypertriglyceridemia-associated missense variants in the polypeptide chain. We hypothesized that regions of functional significance in lipase maturation would be marked by the presence of pathogenic variants.

**Figure 4.**
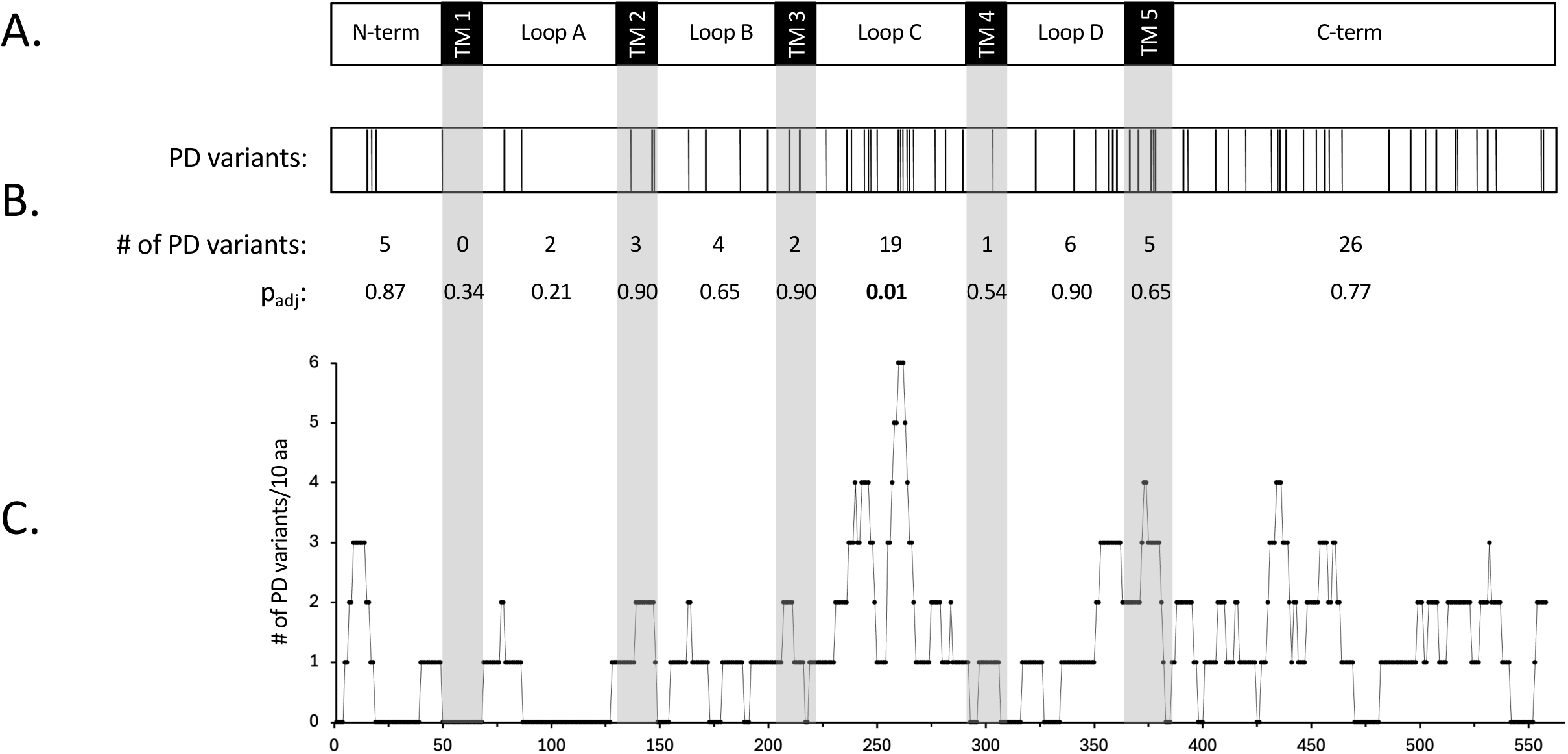
Domain distribution of hTG-associated LMF1 variants. (**A**) Schematic representation of LMF1 domain structure. TM, transmembrane region. (**B**) The top panel shows the domain distribution of 73 hTG-associated missense variants predicted to be detrimental (PD) to LMF1 function. The number of variants and the statistical significance (p_adj_) of enrichment in each domain are shown. (**C**) Graphical representation of variant distribution showing the number of PD variants in a moving window of 10 amino acids.

We used 26 functionally tested LMF1 variants to benchmark a diverse set of variant-effect prediction algorithms (AlphaMissense, ESM1b, PolyPhen2, PROVEAN, REVEL, SIFT) for discrimination between pathogenic and neutral LMF1 variants (**Suppl. Table 1**). Of the predictors tested, PROVEAN exhibited the highest predictive power based on both Matthews correlation coefficient (MCC=0.874) and Receiver Operating Characteristic (AUC=0.977) analysis (**Suppl. Table 2**). In a curated list of 97 hypertriglyceridemia-associated LMF1 variants, PROVEAN predicted 73 as deleterious (**Suppl. Table 1**). While predicted deleterious (PD) variants occur in every soluble and membrane-spanning region except TM1, they exhibit non-uniform distribution in LMF1 structural domains (**Figure 4C**). Notably, the number of PD variants occurring within the ER lumen-facing Loop C represents >2-fold enrichment (p=0.009) over the number expected by chance (**Figure 4B**) indicating the functional importance of this domain in lipase maturation.

## DISCUSSION

Following its discovery in a naturally occurring mutant mouse strain (i.e. *cld*) with severe chylomicronemia, LMF1 has been established as a canonical gene affected in the corresponding human condition, FCS (3). Although collectively responsible for only a small fraction of cases, multiple biallelic LMF1 mutations have been identified in FCS (1, 9, 16, 24–26). As heterozygous variants in canonical FCS genes also contribute to MCS, a much more common manifestation of severe hTG, the allelic landscape of LMF1 has also been extensively explored in this pathology (4, 7–13). Indeed, a curated list of MCS-associated LMF1 alleles currently contains ∼100 variants, most of which remain functionally uncharacterized and their clinical significance uncertain (**Supplemental Table 1**).

To facilitate the functional characterization of LMF1 variants, we previously reported an LMF1 activity assay based on complementation and the reconstitution of LPL maturation in Lmf1-deficient *cld*-mutant cells. This ‘first-generation assay’ has been successfully used to demonstrate the functional impact of LMF1 mutations with large effects (nonsense, frame-shifting) that are typically observed in FCS (1, 16, 17). However, most variants identified in MCS are missense and result in a broad range of allelic effect sizes. Based on our experience with the first-generation assay, it is not well-suited for the identification of low-impact functional variants due to limited sensitivity, precision and throughput. The aim of the present work was to address these limitations and develop an improved LMF1 activity assay.

Previous functional studies of LMF1 variants relied on LPL activity in culture media as the sole measure of cellular lipase maturation (1, 11, 13, 15, 16). Although LPL activity is a proxy for overall cellular lipase maturation activity, it is not only affected by LMF1 function, but also by changes in LMF1 expression under the limiting conditions of the assay. Consequently, earlier assays do not allow discrimination between a variant’s effect on LMF1 activity and its expression. Thus, a principal innovation of the second-generation assay is the assessment of LMF1 protein expression and the use of LMF1 specific activity (i.e. LPL activity/LMF1 protein mass) as the true measure of a variant’s impact on LMF1 function (**Fig. 1**). We achieved this by utilizing GLuc-LMF1 test constructs, which allows the simple and quantitative assessment of LMF1 protein mass by measuring luciferase activity in cell lysates.

The first-generation assay takes advantage of *cld*-mutant mouse cells for complementation. *Cld* is a naturally occurring mutation caused by a retroviral integration in intron 7 of the Lmf1 gene and the mutant allele produces a stable transcript coding for >60% of the full-length protein (1). While the C-terminally truncated mutant Lmf1 protein (Lmf1*^cld^*) lacks lipase maturation activity, its ability to interact with LPL is fully retained (MP unpublished observation), as well as two cysteine residues in Loop C thought to be important for the oxidoreductase activity of LMF1 and LPL secretion (2). To eliminate potential complications associated with the endogenously expressed Lmf1*^cld^* protein, we replaced *cld* hepatocytes with CRISPR-generated LMF1-deficient HEK293 cells in the second-generation assay (**Fig. 3A**). While these cells harbor a frame-shifting deletion in exon 4 and may express a truncated form of LMF1 at low level, this mutant protein lacks Loop C and the ability to interact with LPL (MP unpublished observation) thereby limiting the potential for a dominant negative effect. The use of HEK293 cell line instead of mouse hepatocytes provides the additional advantages of higher transfection efficiency and the elimination of species mismatch between mouse cells and human LMF1 test constructs.

We also addressed a problem stemming from the use of SEAP for normalization in the first-generation assay. As a secreted enzyme, SEAP activity is reduced by the unfolded protein response (18), a process triggered by LMF1 overexpression (MP unpublished observation). Therefore, changes in LMF1 expression due to variable transfection efficiency or altered protein turnover may result in variation in SEAP activity making it poorly suited for normalization. Thus, in the second-generation assay SEAP is replaced with FLuc, a cytosolic enzyme whose activity can be conveniently measured in the same aliquot of cell lysate that is used for the determination of GLuc activity (**Fig. 3B**).

Finally, to increase the throughput of the assay, we replaced the multistep, radiolabeled substrate-based LPL activity measurement with a fluorescence-based assay, which can be performed directly in aliquots of cell culture supernatants (**Fig. 2**). This change allows the entire LMF1 assay to be performed in a few hours in multi-well plates by measuring dual luciferase (i.e. GLuc and FLuc) activities in cell lysates and LPL activity in the corresponding cell culture media. In conclusion, we developed a streamlined LMF1 activity assay suitable for the large-scale functional analysis of LMF1 variants.

We evaluated the performance of the new assay by analyzing 23 missense LMF1 variants from a curated list of 96 variants (**Supplemental Table 1**) identified in patients with severe hTG (4, 7–13). For a direct comparison between the second and first-generation assays, we re-analyzed 8 variants that have been previously reported as neutral in functional analyses with the latter (11, 15). Based on total LMF1 activity (i.e. LPL activity in media) and LMF1 specific activity, the second-generation assay identified 5 and 7 as LOF variants, respectively (**Table 1**). The median specific activity of these variants is 72% (range of 50-78%) of the wild-type protein. Taken together, these results suggest that the second-generation assay has improved sensitivity to detect variants of modest effect sizes.

The assessment of specific, in addition to total, LMF1 activity is a hallmark feature of the second-generation assay. Based on the analysis of 23 variants, these two measures of LMF1 activity were similar (≤10% difference) in most (17 of 23) cases, nonetheless several variants exhibited substantially different total and specific activities (**Table 1**). For instance, mean total and specific activities of R230Q were 84% and 72%, respectively, indicating that higher expression of the mutant protein partially compensates for its reduced activity. Hence, relying on total LMF1 activity leads to erroneous functional assignment (i.e. neutral instead of LOF) of this mutation. A contrasting example is T143M, where specific activity (79%) was markedly higher than total activity (43%) due to decreased mutant protein expression. In this case, total LMF1 activity overestimates the true effect of the variant on LMF1 function. Interestingly, T143M has previously been reported as a gain-of-function variant based on the measurement of total LMF1 activity in a different assay (13). Contrasting with our data, these results suggest increased, not decreased T143M protein level, which may reflect the idiosyncrasies of variants’ impact on protein expression in different cellular contexts. As such effects may not be pertinent to human tissues, expression data are not reported in the present study. Collectively, our results highlight the importance of assessing specific activity in the functional characterization of LMF1 variants.

LMF1 is a transmembrane protein with 5 membrane-spanning and 6 soluble domains. Initial insights into structure-function relationships were provided by functional analyses of nonsense mutations associated with severe hTG (1, 13, 15, 16, 26). These studies revealed that even the shortest truncations of LMF1 (i.e. W464X, Y439X) cause near complete loss of lipase maturation activity. While these results identify the C-terminal region as a critical domain for LMF1 function, they also preclude the use of earlier occurring nonsense mutations for functional domain mapping. To gain initial insights into additional structural determinants in LMF1, we leveraged a curated list of 97 missense variants previously reported in severe hTG (**Supplemental Table 1**). The distribution of 73 predicted deleterious variants revealed significant enrichment in Loop C suggesting an important role of this domain in LMF1 function (**Fig. 4B**). Consistent with this prediction, a recent genetic analysis in a patient with MCS identified a homozygous genomic deletion of exon 6 resulting in the in-frame deletion of amino acids 244-299 within Loop C (27). Our analysis demonstrates that this 56-aa region is the most mutation-enriched segment in the LMF1 polypeptide and includes 14 functionally characterized or predicted LOF variants (**Fig. 4C**). Taken together, these results identify this sub-domain as a critical structural determinant of lipase maturation and warrant further studies to investigate the underlying mechanisms.

Heterozygosity for rare LOF variants in one of the canonical FCS genes has been recognized as a contributor to polygenic risk for severe hTG in a substantial minority (15-25%) of the patient population (28). Whereas most (60-80%) such variants occur in the LPL gene (29), we previously detected increased genetic burden in LMF1 in patients with hTG suggesting that LMF1 variants may also contribute to polygenic risk for this phenotype (30). Although numerous rare LMF1 variants have since been identified in hTG cohorts, the functional significance of most remains unknown (**Supplemental Table 1**). In the present study, we gained initial insights into the allelic effect sizes of hTG-associated LMF1 variants. The analysis of 14 previously uncharacterized missense variants revealed 12 LOF variants with a median LMF1 specific activity of 57% (range of 4-88%) (**Table 1**). The relatively modest allelic effects on LPL activity suggest that in patients with rare heterozygous LMF1 mutations, severe hTG may be the result of significant contributions from TG-raising common variants and non-genetic factors. The LMF1 activity assay developed in the present study will allow a more comprehensive evaluation of hTG-associated rare variants and facilitate further insights into the genetic architecture of severe hTG.

## Data availability

All data described are contained in the article.

## Supplemental data

This article contains supplemental data.

## Acknowledgement

We thank David Stufflebeam for technical assistance.

## Author contributions

Candy Bedoya: Data curation, Formal analysis, Investigation, Methodology, Writing – review & editing. Parmis Rejali, Xanden McCleary, Zachary Hall, Laurie Green, Philip H. Frost: Investigation, Methodology, Writing – review & editing. Michael M. Hoffmann, Mary J. Malloy, Clive Pullinger, John P. Kane: Conceptualization, Data curation, Funding acquisition, Project administration, Supervision, Writing – review & editing. Miklos Peterfy: Conceptualization, Data curation, Funding acquisition, Project administration, Supervision, Visualization, Writing – original draft.

## Notes

Funding: This work was supported by the National Institutes of Health (R01HL127155 and R15HL154071 to M.P.), the Joseph Drown Foundation, the Campini Foundation, and by gifts from Peter Read, Harold Dittmer, Susan Boeing and Donald Yellon.

### Competing Interest Statement

The authors have declared no competing interest.

