## Supplemental data for "Functional characterization of hypertriglyceridemia-associated mutations in Lipase Maturation Factor 1 by a second-generation activity assay"

**Supplemental Table 1.** HTG-associated LMF1 missense variants

| # | Variant | Allele frequency (gnomAD) | Zygosity | HGVS c. | HGVS p. | dbSNP | Reference | Franklin ACMG classification[a] | 2nd-gen functional assay result[b] | LMF1 domain | ESM1b[c] | AlphaMissense[d] | PolyPhen2[e] | PROVEAN[f] | SIFT[g] | REVEL[h] |
|---|---|---|---|---|---|---|---|---|---|---|---|---|---|---|---|---|
| 1 | E13A | nr | HET | 38A>C | Glu13Ala | nr | Puerto-Baracaldo et al. (23) | VUS | | N-term | -3.957 | 0.070 | 0.001 | -0.500 | 0.260 | 0.034 |
| 2 | S14W | 6.45E-07 | cHET | 41C>A | Ser14Trp | rs759977380 | Heidemann et al. (15) | VUS | | N-term | -8.947 | 0.136 | **0.988** | **-1.973** | **0.001** | 0.091 |
| 3 | R16W | 2.56E-06 | HET | 46A>T | Arg16Trp | rs1264860590 | Puerto-Baracaldo et al. (23) | VUS | | N-term | **-11.954** | **0.333** | **0.997** | **-3.121** | **0.002** | **0.165** |
| 4 | R16S | nr | HET | 48G>T | Arg16Ser | nr | Puerto-Baracaldo et al. (23) | VUS | | N-term | **-10.831** | **0.664** | **0.816** | **-2.067** | **0.009** | 0.085 |
| 5 | R18G | 2.76E-05 | HET | 51G>A | Arg18Gly | rs766688892 | Puerto-Baracaldo et al. (23) | VUS | | N-term | **-11.619** | **0.247** | **0.915** | **-2.081** | **0.005** | 0.102 |
| 6 | R35H | 1.91E-05 | HET | 104G>A | Arg35His | rs759097865 | Bashir et al. (3) | VUS | | N-term | -5.858 | 0.090 | 0.003 | -0.540 | **0.121** | 0.025 |
| 7 | G36D | 9.43E-02 | HOM | 107G>A | Gly36Asp | rs111980103 | Johansen et al. (16) | B | | N-term | -4.628 | 0.052 | 0.000 | 0.680 | 0.882 | 0.071 |
| 8 | A38T | 3.83E-05 | HET | 112G>A | Ala38Thr | rs531928966 | Bashir et al. (3) | VUS | | N-term | -5.580 | 0.077 | 0.002 | -0.420 | **0.179** | 0.016 |
| 9 | F49V | 1.29E-06 | HET | 145T>G | Phe49Val | rs965954981 | Abedi et al. (1) | VUS | | N-term | **-12.065** | **0.534** | **0.842** | **-3.706** | **0.004** | **0.236** |
| 10 | A59V | 1.30E-04 | HET | 176C>T | Ala59Val | rs759181295 | Mendes et al. (18) | LB | | TM1 | -8.264 | 0.122 | 0.704 | -1.628 | 0.228 | **0.149** |
| 11 | L78P | 2.49E-06 | HOM | 233T>C | Leu78Pro | rs759535337 | Garay-Garcia et al. (11) | LP | | Loop A | **-14.819** | **0.953** | **1.000** | **-5.151** | **0.000** | **0.826** |
| 12 | P86L | 1.99E-05 | cHET | 257C>T | Pro86Leu | rs1269525472 | Liu et al. (17) | VUS | | Loop A | **-10.870** | **0.774** | **0.998** | **-3.250** | **0.003** | **0.713** |
| 13 | Q96E | nr | HET | 286C>G | Gln96Glu | nr | Puerto-Baracaldo et al. (23) | VUS | | Loop A | -3.878 | 0.075 | 0.000 | -0.451 | 1.000 | 0.037 |
| 14 | D100N | 2.29E-05 | HET | 298G>A | Asp100Asn | rs35124265 | Gill et al. (12) | VUS | | Loop A | -5.694 | 0.070 | 0.008 | 0.040 | **0.145** | 0.108 |
| 15 | R101T | 8.34E-04 | HET | 302G>C | Arg101Thr | rs147688306 | Johansen et al. (16) | B | | Loop A | -8.909 | 0.120 | 0.028 | -1.390 | 0.565 | 0.061 |
| 16 | M122V | 4.96E-06 | HET | 364A>G | Met122Val | rs1207954762 | Johansen et al. (16) | VUS | | Loop A | -7.565 | 0.085 | 0.001 | -1.170 | 0.689 | **0.229** |
| 17 | L128F | 5.96E-04 | cHET | 384G>C | Leu128Phe | rs115313199 | present study | B | | TM2 | -5.909 | 0.067 | 0.551 | -0.830 | 0.712 | **0.159** |
| 18 | S137L | 1.80E-05 | cHET | 410C>T | Ser137Leu | rs549008037 | Peterfy et al. (20) | LP | pLOF | TM2 | **-14.963** | **0.634** | **1.000** | **-3.250** | **0.046** | **0.770** |
| 19 | S138C | 2.13E-04 | HET | 413C>G | Ser138Cys | rs200382562 | Dancer et al. (25) | LB | | TM2 | -7.166 | 0.068 | **0.980** | -1.156 | **0.170** | **0.347** |
| 20 | T143M | 8.99E-05 | HET | 428C>T | Thr143Met | rs375529211 | Dancer et al. (25) | LB | Normal | TM2 | -6.591 | 0.107 | 0.773 | -0.830 | **0.144** | 0.106 |
| 21 | N147K | 2.67E-05 | HOM | 441C>G | Asn147Lys | rs182685983 | Bedoya et al. (30) | LP | pLOF | TM2 | **-13.584** | **0.976** | **0.992** | **-4.480** | **0.003** | **0.222** |
| 22 | M148K | nr | HET | 443T>A | Met148Lys | nr | Blokhina et al. (32) | VUS | | TM2 | **-14.253** | **0.667** | 0.473 | **-3.690** | **0.175** | **0.261** |
| 23 | M159V | 1.46E-04 | HET | 475A>G | Met159Val | rs142481016 | present study | B | | Loop B | -8.934 | 0.106 | 0.064 | -1.470 | 0.268 | 0.124 |
| 24 | V164A | 4.01E-03 | HET | 491T>C | Val164Ala | rs35663121 | Puerto-Baracaldo et al. (23) | B | | Loop B | -5.722 | 0.149 | 0.714 | **-2.760** | **0.132** | **0.228** |
| 25 | G172R | 4.63E-05 | HOM | 514G>A | Gly172Arg | rs201406396 | Dancer et al. (25) | LP | | Loop B | **-14.680** | **0.871** | **0.999** | **-5.727** | 0.310 | **0.307** |
| 26 | C188Y | 3.10E-06 | cHET | 563G>A | Cys188Tyr | rs2548363369 | present study | VUS | | Loop B | **-15.499** | **0.952** | **0.984** | **-6.060** | **0.004** | **0.330** |
| 27 | P201L | 1.67E-05 | HET | 602C>T | Pro201Leu | rs565159724 | Gill et al. (12) | VUS | | Loop B | **-12.350** | **0.312** | **1.000** | **-6.890** | **0.000** | **0.358** |
| 28 | R211Q | 6.70E-05 | HET | 632G>C | Arg211Pro | rs182946890 | Gill et al. (12) | VUS | | TM3 | -7.819 | **0.191** | **0.994** | **-2.640** | **0.008** | **0.379** |
| 29 | R216T | nr | HET | 647G>C | Arg216Thr | nr | Dong et al. (9) | VUS | pLOF | TM3 | **-13.659** | **0.991** | **0.991** | **-4.590** | **0.004** | **0.449** |
| 30 | G228A | 6.44E-07 | HET | 683G>C | Gly228Ala | rs754772870 | Gill et al. (12) | VUS | pLOF | Loop C | **-9.595** | **0.375** | **0.975** | **-4.710** | **0.131** | **0.219** |
| 31 | G228E | 6.01E-04 | HET | 683G>A | Gly228Glu | rs754772870 | Dron et al. (10) | VUS | pLOF | Loop C | **-12.527** | **0.600** | **0.884** | **-6.470** | **0.024** | **0.198** |
| 32 | G228V | 1.29E-05 | HET | 683G>T | Gly228Val | rs754772870 | Dancer et al. (25) | VUS | | Loop C | **-11.028** | **0.721** | **0.998** | **-7.392** | **0.030** | **0.249** |
| 33 | R230Q | 1.48E-03 | HET | 689G>A | Arg230Gln | rs192224688 | Surendran et al. (26) | B | pLOF | Loop C | -1.905 | 0.068 | 0.027 | -0.110 | 0.764 | **0.147** |
| 34 | M238T | 1.04E-04 | HET | 713T>C | Met238Thr | rs370864352 | Dron et al. (10) | VUS | pLOF | Loop C | **-11.209** | **0.480** | **0.999** | **-5.070** | **0.001** | **0.492** |
| 35 | F240L | 6.45E-07 | HET | 718T>C | Phe240Leu | rs34295987 | Gill et al. (12) | LB | | Loop C | -9.242 | **0.907** | 0.367 | **-3.910** | **0.009** | **0.181** |
| 36 | P246R | 1.26E-06 | HOM | 737C>G | Pro246Arg | rs1307363051 | Bedoya et al. (30) | VUS | pLOF | Loop C | **-16.998** | **0.871** | **1.000** | **-8.460** | **0.000** | **0.685** |
| 37 | P248S | nr | HOM | 742C>T | Pro248Ser | nr | Hegele et al. (14) | VUS | | Loop C | **-14.514** | **0.811** | **1.000** | **-7.516** | **0.000** | **0.796** |
| 38 | P248T | nr | HET | 742C>A | Pro248Thr | rs2544528154 | Puerto-Baracaldo et al. (23) | VUS | | Loop C | **-16.806** | **0.760** | **1.000** | **-7.516** | **0.000** | **0.793** |
| 39 | N249S | 6.28E-06 | HET | 746A>G | Asn249Ser | rs766214202 | Plengpanich et al. (22) | VUS | | Loop C | -5.662 | 0.062 | **0.966** | **-4.260** | **0.007** | **0.271** |
| 40 | A252V | 1.81E-05 | HOM | 755C>T | Ala252Val | rs773491556 | Bashir et al. (3) | VUS | | Loop C | **-14.316** | **0.501** | **0.996** | **-3.030** | **0.007** | **0.437** |
| 41 | F262L | 1.24E-05 | HET | 786C>A | Phe262Leu | rs770348738 | Gill et al. (12) | VUS | | Loop C | -6.474 | **0.413** | 0.117 | **-3.580** | **0.175** | 0.066 |
| 42 | H263Y | 4.60E-05 | HET | 787C>T | His263Tyr | rs746165846 | Dron et al. (10) | VUS | | Loop C | **-13.589** | **0.583** | **1.000** | **-5.204** | **0.020** | **0.583** |
| 43 | R264C | 4.97E-06 | HET | 790C>T | Arg264Cys | rs777579889 | Surendran et al. | VUS | pLOF | Loop C | **-9.538** | 0.101 | **0.999** | **-5.240** | **0.003** | **0.490** |
| 44 | E266K | 1.49E-05 | HET | 796G>A | Glu266Lys | rs778529081 | Mendes et al. (18) | VUS | | Loop C | **-11.661** | **0.685** | **1.000** | **-3.091** | **0.032** | **0.448** |
| 45 | T267M | 8.07E-05 | HET | 800C>T | Thr267Met | rs754428234 | Mendes et al. (18) | VUS | | Loop C | -6.479 | 0.069 | **0.984** | **-2.030** | **0.182** | **0.358** |
| 46 | S269C | nr | HET | 805A>T | Ser269Cys | nr | Bashir et al. (3) | VUS | | Loop C | **-10.480** | **0.204** | **1.000** | **-3.837** | **0.011** | **0.216** |
| 47 | I273L | 3.53E-05 | HET | 817A>C | Ile273Leu | rs762056405 | Dron et al. (10) | VUS | | Loop C | -7.542 | 0.067 | 0.022 | -1.340 | 0.230 | 0.100 |
| 48 | F279L | 4.70E-03 | HET | 837C>A | Phe279Leu | rs61745065 | Puerto-Baracaldo et al. (23) | B | | Loop C | -6.984 | **0.667** | 0.107 | **-4.237** | **0.086** | **0.155** |
| 49 | G284S | 4.96E-05 | HET | 850G>A | Gly284Ser | rs928471580 | Dron et al. (10) | VUS | pLOF | Loop C | -7.709 | **0.195** | **0.999** | **-1.870** | 0.705 | **0.302** |
| 50 | A287V | 3.53E-05 | HET | 860C>T | Ala287Val | rs755996576 | Plengpanich et al. (22) | VUS | | Loop C | -4.595 | 0.085 | 0.637 | 0.530 | 0.512 | **0.185** |
| 51 | G292R | 1.36E-05 | HOM | 874G>C | Gly292Arg | rs764154907 | Abedi et al. (1) | VUS | | Loop C | **-9.704** | **0.969** | **1.000** | **-7.283** | 0.200 | **0.612** |
| 52 | G306R | 2.54E-05 | HET | 916G>A | Gly306Arg | rs372159961 | Dron et al. (10) | VUS | | TM4 | **-14.194** | **0.867** | **1.000** | **-7.620** | **0.000** | **0.694** |
| 53 | M316V | 5.58E-06 | HET | 946A>G | Met316Val | rs765861000 | Gill et al. (12) | VUS | | TM4 | -6.780 | 0.058 | 0.002 | 0.991 | 0.410 | **0.148** |
| 54 | A326T | 4.34E-05 | HET | 976G>A | Ala326Thr | rs769766473 | Johansen et al. (16) | VUS | | Loop D | -8.230 | 0.084 | **0.978** | **-2.010** | 0.333 | 0.095 |
| 55 | L344P | 6.21E-07 | HET | 1031T>C | Leu344Pro | rs2069779961 | present study | VUS | | Loop D | -4.083 | **0.432** | 0.036 | **-2.510** | **0.148** | **0.270** |
| 56 | M346V | 1.00E-04 | HET | 1036A>G | Met346Val | rs201767825 | Plengpanich et al. (22) | LB | | Loop D | -2.528 | 0.114 | 0.001 | -0.220 | 0.237 | 0.096 |

| | | | | | | | | | | | | | | | | |
|---|---|---|---|---|---|---|---|---|---|---|---|---|---|---|---|---|
| 57 | R351Q | 1.76E-03 | HET | 1052G>A | Arg351Gln | rs192520307 | Puerto-Baracaldo et al. (23) | B | Normal | Loop D | -4.218 | 0.069 | 0.004 | 0.150 | 0.748 | 0.012 |
| 58 | R354W | 1.65E-02 | HOM | 1060C>T | Arg354Trp | rs143076454 | Surendran et al. (26) | B | pLOF | Loop D | -5.884 | 0.081 | 0.002 | **-2.760** | **0.085** | 0.015 |
| 59 | R354Q | 8.99E-04 | HET | 1060C>G | Arg354Gly | rs138461953 | Puerto-Baracaldo et al. (23) | B | | Loop D | -3.875 | 0.068 | 0.001 | -0.090 | 0.833 | 0.016 |
| 60 | G360S | 1.33E-05 | HET | 1078G>A | Gly360Ser | rs762677583 | Gill et al. (12) | VUS | pLOF | Loop D | -2.818 | 0.106 | **0.949** | **-3.100** | **0.097** | 0.082 |
| 61 | V362G | 9.46E-05 | HET | 1085T>G | Val362Gly | rs368408082 | Abedi et al. (1) | VUS | | Loop D | -4.822 | 0.090 | 0.430 | **-2.477** | 0.278 | **0.176** |
| 62 | R364Q | 2.68E-02 | HOM | 1091G>A | Arg364Gln | rs35168378 | Surendran et al. (26) | B | pLOF | Loop D | -7.301 | **0.195** | **0.977** | **-2.650** | **0.080** | **0.179** |
| 63 | S370L | 3.35E-05 | HET | 1109C>T | Ser370Leu | rs769234511 | D'Erasmo et al. (8) | VUS | | TM5 | -0.089 | 0.084 | 0.209 | **-2.733** | 0.722 | 0.091 |
| 64 | L374Q | 3.83E-04 | HET | 1121T>A | Leu374Gln | rs778846872 | Abedi et al. (1) | VUS | | TM5 | **-17.419** | **0.893** | **0.999** | **-5.233** | **0.002** | **0.523** |
| 65 | V380M | 2.91E-04 | HET | 1138G>A | Val380Met | rs201734228 | Mendes et al. (18) | VUS | | TM5 | -8.107 | 0.096 | 0.672 | **-2.006** | **0.005** | **0.165** |
| 66 | P381L | 6.20E-07 | cHET | 1142C>T | Pro381Leu | rs1408203743 | Montalvo et al. (19) | VUS | | TM5 | **-11.017** | **0.552** | **0.883** | **-7.124** | **0.003** | **0.427** |
| 67 | V382M | 7.01E-05 | HET | 1144G>A | Val382Met | rs370036895 | Dron et al. (10) | VUS | pLOF | TM5 | **-12.004** | **0.566** | **1.000** | **-2.160** | **0.009** | **0.327** |
| 68 | T395I | 1.46E-04 | cHET | 1184C>T | Thr395Ile | rs186694298 | Liu et al. (17) | LB | | C-term | **-11.325** | **0.207** | 0.591 | **-5.510** | **0.013** | **0.167** |
| 69 | F397I | 6.20E-07 | HET | 1189T>A | Phe397Ile | rs2544495863 | Mendes et al. (18) | VUS | | C-term | **-11.672** | **0.837** | 0.642 | **-5.459** | **0.005** | **0.497** |
| 70 | G410R | 4.65E-04 | HET | 1228G>A | Gly410Arg | rs199713950 | Dancer et al. (25) | B | | C-term | **-13.950** | **0.818** | **0.989** | **-7.290** | **0.000** | **0.466** |
| 71 | R416Q | 1.93E-05 | HOM | 1247G>A | Arg416Gln | rs762640109 | Cao et al. (4) | VUS | | C-term | **-10.212** | **0.479** | **0.998** | **-3.680** | **0.000** | **0.761** |
| 72 | T424R | 3.41E-05 | HET | 1271C>G | Thr424Arg | rs370807438 | present study | VUS | | C-term | **-15.399** | **0.433** | **0.996** | **-5.050** | **0.001** | **0.614** |
| 73 | A431D | 1.36E-03 | HET | 1292C>A | Ala431Asp | rs115416993 | Dancer et al. (25) | B | Normal | C-term | -8.872 | 0.079 | 0.001 | 0.720 | 0.617 | 0.026 |
| 74 | D433N | 7.69E-05 | HET | 1297G>A | Asp433Asn | rs201927375 | Plengpanich et al. (22) | LB | | C-term | -8.732 | 0.081 | 0.076 | 0.720 | 0.318 | 0.052 |
| 75 | W436C | nr | cHET | 1308G>C | Trp436Cys | nr | Gong et al. (13) | VUS | | C-term | **-16.232** | **0.981** | **1.000** | **-11.781** | **0.000** | **0.600** |
| 76 | Y439C | 4.66E-05 | HOM | 1316A>G | Tyr439Cys | rs764885027 | Blokhina et al. (32) | VUS | | C-term | **-11.768** | **0.470** | **1.000** | **-8.180** | **0.000** | **0.403** |
| 77 | E440K | 3.97E-05 | HET | 1318G>A | Glu440Lys | rs778629426 | present study | VUS | | C-term | **-11.506** | **0.181** | **0.997** | **-3.150** | **0.015** | **0.354** |
| 78 | C443Y | nr | HET | 1328G>A | Cys443Tyr | nr | Bashir et al. (3) | VUS | | C-term | **-11.251** | **0.709** | 0.083 | **-7.410** | **0.001** | **0.364** |
| 79 | R451W | 3.19E-03 | HET | 1351C>T | Arg451Trp | rs138205062 | Johansen et al. (16) | B | pLOF | C-term | **-9.475** | 0.111 | **1.000** | **-6.240** | **0.001** | **0.197** |
| 80 | P457L | 6.94E-05 | cHET | 1370C>T | Pro457Leu | rs757628838 | present study | VUS | pLOF | C-term | **-12.382** | **0.654** | **1.000** | **-9.190** | **0.000** | **0.762** |
| 81 | R461H | 4.03E-05 | HET | 1382G>A | Arg461His | rs557053661 | Dancer et al. (25) | VUS | | C-term | **-11.343** | **0.587** | **1.000** | **-4.529** | **0.000** | **0.685** |
| 82 | R461C | 1.49E-05 | HET | 1381C>T | Arg461Cys | rs753409067 | Abedi et al. (1) | VUS | LOF | C-term | **-11.758** | **0.702** | **1.000** | **-7.252** | **0.000** | **0.794** |
| 83 | D463N | 1.24E-05 | HET | 1387G>A | Asp463Asn | rs750391160 | Dron et al. (10) | VUS | | C-term | **-14.092** | **0.839** | **1.000** | **-4.595** | **0.000** | **0.539** |
| 84 | A469T | 4.83E-04 | HET | 1405G>A | Ala469Thr | rs181731943 | Johansen et al. (16) | B | | C-term | **-9.746** | **0.269** | **1.000** | **-3.370** | **0.010** | **0.343** |
| 85 | D491N | 1.05E-04 | HET | 1471G>A | Asp491Asn | rs532127028 | Plengpanich et al. (22) | LB | | C-term | -5.271 | **0.183** | **0.920** | **-3.370** | **0.008** | **0.264** |
| 86 | N501Y | 1.24E-06 | HET | 1501A>T | Asn501Tyr | rs774296427 | Plengpanich et al. (22) | VUS | | C-term | **-14.105** | **0.360** | **0.991** | **-7.110** | **0.001** | **0.432** |
| 87 | A504V | 5.21E-05 | HET | 1511C>T | Ala504Val | rs369478194 | Plengpanich et al. (22) | LB | | C-term | -4.470 | 0.085 | 0.110 | -1.120 | 0.255 | 0.095 |
| 88 | P508L | 1.81E-05 | HET | 1523C>T | Pro508Leu | rs372213215 | Guo et al. (31) | VUS | | C-term | -7.612 | 0.087 | **1.000** | **-6.200** | **0.000** | **0.296** |
| 89 | R513Q | 3.33E-05 | HET | 1538G>A | Arg513Gln | rs748287562 | present study | VUS | pLOF | C-term | **-9.629** | **0.240** | **1.000** | **-3.560** | **0.005** | **0.474** |
| 90 | S522I | nr | HET | 1565G>T | Ser522Ile | nr | Johansen et al. (16) | VUS | pLOF | C-term | **-11.093** | **0.623** | **0.986** | **-3.360** | **0.011** | **0.311** |
| 91 | R523H | 1.43E-03 | HOM | 1568G>A | Arg523His | rs151137164 | Surendran et al. (26) | B | pLOF | C-term | -7.573 | 0.075 | 0.359 | **-2.410** | **0.112** | **0.343** |
| 92 | E531D | 9.21E-04 | HET | 1593G>C | Glu531Asp | rs190958016 | Gill et al. (12) | B | Normal | C-term | -4.457 | 0.089 | 0.000 | -0.860 | 0.482 | **0.415** |
| 93 | G532S | 1.87E-06 | HET | 1594G>A | Gly532Ser | rs1315804529 | Gill et al. (12) | VUS | pLOF | C-term | -9.301 | **0.192** | **1.000** | **-4.560** | **0.007** | **0.432** |
| 94 | R537W | 5.04E-05 | HOM | 1609C>T | Arg537Trp | rs555435528 | Dancer et al. (25) | VUS | | C-term | **-12.858** | **0.421** | **1.000** | **-7.180** | **0.001** | 0.095 |
| 95 | G541R | 1.10E-04 | HOM | 1621G>A | Gly541Arg | rs377058908 | Bashir et al. (3) | B | | C-term | **-9.748** | **0.260** | **0.998** | **-6.610** | **0.004** | **0.302** |
| 96 | P562R | 2.00E-02 | HOM | 1685C>G | Pro562Arg | rs4984948 | Surendran et al. (26) | B | pLOF | C-term | -4.256 | 0.104 | 0.754 | **-2.830** | **0.084** | **0.292** |
| 97 | L563R | 7.96E-05 | HET | 1688T>G | Leu563Arg | rs199591347 | Plengpanich et al. (22) | LB | | C-term | -1.337 | 0.090 | 0.500 | **-1.870** | 0.217 | **0.474** |

nr, not reported
[a], B, benign; LB, likely benign; LP, likely pathogenic; P, pathogenic; VUS, variant of unknown significance
[b], Functional assessment is based on LMF1 specific activity. Normal: >90% activity; pLOF (partial loss-of-function): 10-90% activity; LOF: <10% activity
[c], ESM1b scores ≤ -9.459 are indicated in bold.
[d], AlphaMissense scores ≥ 0.179 are indicated in bold.
[e], PolyPhen2 scores ≥ 0.8 are indicated in bold.
[f], PROVEAN scores ≤ -1.807 are indicated in bold.
[g], SIFT scores ≤ 0.191 are indicated in bold.
[h], REVEL scores ≥ 0.141 are indicated in bold.

**Supplemental Table 2.** Comparison of variant effect predictors based on ROC and MCC analysis

| | | | | | | MCC analysis | | | | | | | | | | |
|---|---|---|---|---|---|---|---|---|---|---|---|---|---|---|---|---|
| **Predictor** | **Direction** | **N used** | **Positives** | **Negatives** | **ROC-AUC** | **Best threshold** | **Positive call rule** | **MCC** | **Accuracy** | **Precision** | **Sensitivity** | **F1** | **TP** | **TN** | **FP** | **FN** |
| PROVEAN | lower | 26 | 22 | 4 | 0.977 | -1.807 | PROVEAN ≤ -1.80705 | 0.874 | 0.962 | 1.000 | 0.955 | 0.977 | 21 | 4 | 0 | 1 |
| PolyPhen2 | higher | 26 | 22 | 4 | 0.943 | 0.800 | PolyPhen2 ≥ 0.8 | 0.640 | 0.846 | 1.000 | 0.818 | 0.900 | 18 | 4 | 0 | 4 |
| SIFT | lower | 26 | 22 | 4 | 0.920 | 0.191 | SIFT ≤ 0.191 | 0.603 | 0.885 | 0.952 | 0.909 | 0.930 | 20 | 3 | 1 | 2 |
| AlphaMissense | higher | 26 | 22 | 4 | 0.864 | 0.179 | AM ≥ 0.17876 | 0.498 | 0.731 | 1.000 | 0.682 | 0.811 | 15 | 4 | 0 | 7 |
| ESM1b | lower | 26 | 22 | 4 | 0.818 | -9.459 | ESM1b ≤ -9.45905 | 0.461 | 0.692 | 1.000 | 0.636 | 0.778 | 14 | 4 | 0 | 8 |
| REVEL | higher | 26 | 22 | 4 | 0.818 | 0.141 | REVEL ≥ 0.14103 | 0.603 | 0.885 | 0.952 | 0.909 | 0.930 | 20 | 3 | 1 | 2 |

Abreviations: ROC, Receiver Operating Characteristic ; MCC, Matthews correlation coefficient; AUC, area under the curve; TP, true positives; TN, true negatives; FP, false positives; FN, false negatives
